# Splicing regulation and intron evolution in the short-intron ciliate model of endosymbiosis *Paramecium bursaria*

**DOI:** 10.1101/2025.08.05.668791

**Authors:** Thi Ngan Giang Nguyen, Kamal Md Mostafa, Chien-Ling Lin, Jun-Yi Leu

## Abstract

The integration of symbionts into host cells during endosymbiosis significantly alters gene expression and cell physiology. Though alternative splicing facilitates cellular adaptation through rapid modulation of gene expression and protein isoform diversity, its regulatory role during endosymbiosis remains poorly understood. *Paramecium bursaria*, which harbors hundreds of *Chlorella variabilis* algae within its cytoplasm, offers a powerful model to study splicing during endosymbiosis, especially given its exceptionally short introns (median ∼24 nt). Using time-course RNA sequencing of symbiotic and aposymbiotic cells, we found that splicing, especially of 5’ proximal introns, enhances gene expression. Moreover, we identified 883 genes with differentially spliced introns, particularly enriched in transmembrane transporters essential for establishing nutrient exchange between a host cell and algal symbionts. Splicing regulation correlated with expression changes in conserved spliceosome components, implicating that these factors act as splicing enhancers or repressors during symbiosis. By exploring intron orthology across ciliates, we found that conserved introns exhibited more efficient splicing, characterized by lower GC content and uniform length, suggesting that intron evolution favors features that optimize expression. Our study reveals how splicing contributes to host adaptation during endosymbiosis and highlights the evolutionary dynamics of short introns in eukaryotes.

**Key points:** *Paramecium* retains conserved core spliceosome genes but employs unique splicing regulation.

Alternative splicing regulates metabolism during endosymbiosis, especially for transporter genes.

Older introns evolve toward ∼24 nt length and low GC for efficient splicing and expression.

## Introduction

### Molecular basis of RNA splicing

RNA splicing is a crucial post-transcriptional process in eukaryotes, involving exon ligation and intron removal. The U2-dependent splicing pathway is the primary mechanism for intron removal in eukaryotic pre-mRNA, which proceeds through stepwise assembly of the spliceosome on splice junctions (Yang, Beutler et al. 2022). The major components of the U2-dependent spliceosome include five essential small nuclear ribonucleoproteins (i.e., the U1, U2, U4/U6, and U5 snRNPs), which are comprised of associated U-rich small nuclear RNAs (UsnRNAs) and other associated proteins. In human cells, the 5’ splice site (5’SS), branch point site (BPS), and 3’ splice site (3’SS) are recognized by U1 snRNP, Splicing Factor 1 (SF1), and U2 Small Nuclear RNA Auxiliary Factor 1 (U2AF1), respectively, together forming the E complex. SF1 is then displaced by U2 snRNP to form the A complex. The U4/U6.U5 tri-snRNP subsequently joins to create the B complex. Rearrangements of the B complex together with displacement of the U1 and U4 snRNPs lead to the formation of the active B complex and then the B* complex, which catalyzes the first step of splicing, resulting in the C complex. Further rearrangements lead to formation of the C* complex, which carries out the second catalytic step by ligating the exons and allowing spliceosome components to be recycled.

### Regulated alternative splicing enriches proteome diversity

Alternative splicing (AS), which allows for various combinations of exon inclusion or exclusion, intron retention, and the use of alternative splice sites, enables a single gene to produce multiple protein isoforms. In humans, over 90% of multi-exon genes undergo alternative splicing, generating diverse proteins that play different roles in a wide range of cellular processes (Pan, Shai et al. 2008). There are five primary types of alternative splicing events: exon skipping, alternative 3’ splice site, alternative 5’ splice site, mutually exclusive exons, and intron retention (Verta and Jacobs 2022). Alternative splicing isoforms enable the cell to regulate transcriptional levels and they increase proteome diversity. When alternative splicing introduces premature termination codons (PTCs), it activates the nonsense-mediated decay (NMD) pathway, leading to mRNA degradation (Kervestin and Jacobson 2012). The NMD pathway has been shown to regulate transcript levels and ensure fidelity of the splicing process (Alonso 2005, Palacios 2013, Miller and Pearce 2014). In contrast, in-frame alternative splicing events that do not generate PTCs result in changes in protein sequences and structures, thereby altering protein activity or cellular localization (Birzele, Csaba et al. 2008, Kjer-Hansen and Weatheritt 2023). For example, alternative splicing of exons 6 and 7 in the cell division cycle protein 42 (CDC42) affects its lipid modification, thereby influencing its distribution in neuronal cell compartments (Lee, Zdradzinski et al. 2021). In the case of the apoptotic protein Caspase-9, exon skipping generates a short isoform, Caspase-9S, which competes with the long isoform to inhibit apoptosis (Seol and Billiar 1999).

Alternative splicing is a crucial aspect of the regulation of various biological processes, including cell identity and circadian rhythms (Nilsen and Graveley 2010, Kalsotra and Cooper 2011, McGlincy, Valomon et al. 2012, Baralle and Giudice 2017). It plays a significant role in driving cell differentiation, with splicing patterns varying widely among tissues and cell types (Olivieri, Dehghannasiri et al. 2021, Zhang, Guo et al. 2025). For example, in humans, the *+a-b+c* isoform of Ribosomal Protein S24 (RPS24) is the most dominant in epithelial cells, whereas it is barely detectable in non-epithelial cells (Olivieri, Dehghannasiri et al. 2021). Similarly, Mitogen-activated Protein Kinase 7 (MAP3K7) isoform usage shifts from a short isoform predominant in undifferentiated progenitor cells to a long isoform dominant in mature epidermal keratinocytes (Takashima, Sun et al. 2024).

### Host adaptation during endosymbiosis through alternative splicing

The process of endosymbiosis involves the integration of symbionts into host cells, forming a mutually beneficial relationship. In this interaction, endosymbionts confer novel phenotypic traits on the host, enabling its exploitation of more diverse environmental resources. In return, the symbionts rely on the host for essential nutrients (Archibald 2015). One of the most well-known and ancient examples of such a relationship is the “Endosymbiosis Theory”, which explains the origin of mitochondria and chloroplasts through integration of ancestral prokaryotes into early eukaryotic cells (Archibald 2015, Martin, Garg et al. 2015). This integrative process can fundamentally alter host cell biology, such as by transforming photosynthetic cyanobacteria into chloroplasts within plant cells or driving the development of specialized organelles to accommodate endosymbionts (Archibald 2015, von der Dunk, Hogeweg et al. 2023). To maintain such symbiotic relationships, both host cells and endosymbionts must undergo extensive physiological and genomic adaptations, including horizontal gene transfer (HGT) and the evolution of mechanisms for nutrient and resource exchange (Keeling 2013, Martin, Garg et al. 2015, Wilson and Duncan 2015, Kelly 2021, Bennett, Kwak et al. 2024). Although previous research has demonstrated gene family expansion and shifts in gene expression related to host stress responses and metabolism arising from endosymbiosis (Kelly, Carlson et al. 2025), the contribution of alternative splicing to these adaptive processes is not well understood. Given its role in rapidly modulating gene expression and expanding proteomic complexity, alternative splicing may be a key, yet underexplored, mechanism contributing to host adaptation during symbiotic integration.

### RNA splicing regulation in Paramecium

*Paramecium bursaria* serves as an excellent model for studying endosymbiosis. *P. bursaria* stably hosts hundreds of algal cells in its cytoplasm, and these endosymbiotic algae are shielded from lysosome digestion by the perialgal vacuole (PV) beneath the host cytoplasmic membrane (Fujishima and Kodama 2012, He, Wang et al. 2019, Jenkins 2024). *P. bursaria* symbiotic cells typically display increased size and growth rate, and reduced numbers of mitochondria underneath the cell membrane where the PV is located (He, Wang et al. 2019, Kodama and Fujishima 2022). A differential gene expression analysis between symbiotic and aposymbiotic *P. bursaria* cells revealed downregulation of redox, aminotransferase and ribosomal proteins in host cells and upregulation of Hsp70, the Myb transcription factor and histidine kinase pathway genes (Yuuki Kodama 2014). Even though various studies have been conducted on transcriptional regulation in the context of endosymbiosis (Yuuki Kodama 2014, Suzuki, Ishida et al. 2016, Ferrarini, Vallier et al. 2023, Abresch, Bell et al. 2024), the functional role of alternative splicing regulation in endosymbiosis remains unexplored.

Given its importance, RNA splicing is tightly regulated by cis-acting regulatory sequences and trans-acting splicing factors (Hang, Wan et al. 2015, Plaschka, Lin et al. 2018, Chao, Jiang et al. 2021). Cis-acting elements include exonic splicing enhancers, intronic splicing enhancers, exonic splicing silencers, and intronic splicing silencers. Enhancer elements are typically bound by splicing activators such as serine/arginine-rich (SR) proteins, which promote splicing. In contrast, silencer elements are bound by repressor proteins such as heterogeneous nuclear ribonucleoproteins (hnRNPs), leading to splicing inhibition (Wang, Xiao et al. 2006, Wang and Burge 2008, Wang, Liu et al. 2015). Since alternative splicing is regulated in a cell-type-specific manner (Boutz, Stoilov et al. 2007, Yang, Hung et al. 2014, Olivieri, Dehghannasiri et al. 2021, Zhang, Guo et al. 2025), investigating the relationship between splicing factors and splicing efficiency in *P. bursaria* may help uncover key regulators involved in the endosymbiosis process.

### Evolution of extremely short introns in ciliates

Introns exert a crucial role in various evolutionary processes due to their links to regulating adaptive gene expression. Intron characteristics vary across species, with lengths ranging from 15 nucleotides in the ciliate *Stentor coeruleus* to over 1 million nucleotides in humans (Yu, Yang et al. 2002, Nuadthaisong, Phetruen et al. 2022). Although most modern vertebrate species exhibit a bimodal distribution of intron lengths (Yu, Yang et al. 2002), ciliate species such as *Paramecium* and *Tetrahymena* are notable for predominantly harboring extremely short introns, with the majority being under 100 nucleotides. This distinct intron architecture makes ciliates excellent models for studying the evolution and regulation of short intron splicing (Bondarenko and Gelfand 2016). Despite having extremely short intronic sequences, intron splicing in ciliates still plays a significant role in gene regulation (Saudemont, Popa et al. 2017, Gnan, Matelot et al. 2022, Ryll, Rothering et al. 2022). Thus, comparative studies of intron evolution across ciliates can offer valuable insights into the evolutionary history and functional adaptation of short introns, as well as their implications for transcriptomic regulation.

In this study, we examined time-course RNA sequencing data from both symbiotic (symbiont-bearing) and aposymbiotic (symbiont-free) *P. bursaria* cells to quantify alternative splicing across stages. Doing so allowed us to uncover the relationship between splicing efficiency and gene expression in *P. bursaria*. Differential analysis of intron retention rates between the symbiotic and aposymbiotic cells revealed specific splicing regulation, particularly in genes associated with transmembrane transporter activity, a function that is closely linked to endosymbiosis-linked nutrient exchange. Furthermore, through comparative intron orthology across *Paramecium* species, we found that intron length and GC content have evolved to optimize splicing efficiency. Collectively, our findings highlight an evolutionary refinement of intron architecture to enhance splicing precision and metabolic fitness in the context of endosymbiotic adaptation.

## Materials and Methods

### *Paramecium bursaria* strain and culture conditions

We used the *P. bursaria* DK2 strain, which harbors the endogenous endosymbiont *Chlorella variabilis*. DK2 is an offspring of the Dd1 and KM2 strains, and details of the process for creating DK2 have been described previously (Cheng, Liu et al. 2020). *P. bursaria* cells were cultured on 2.5 % Boston lettuce in Dryl’s solution medium and fed with *Klebsiella pneumoniae* (NBRC 100048 strain). Cell cultures were maintained at 23 °C with a 12-hour light/dark cycle. Bacteria-containing lettuce media was refreshed every two days. To produce aposymbiotic (white) *Paramecium* strains, we used cycloheximide (10 μg/ml) to treat symbiotic (green) cells, as described previously (Kodama, Inouye et al. 2011).

### Genome annotation file and intron annotation

Like other ciliate species, *P. bursaria* possesses a diploid genome displaying allele duplication. The genome annotation file (GTF) was assembled according to a method described previously (Cheng, Liu et al. 2020). We separated the diploid genome annotation into two haplotypes and identified 14,920 functional genes as representatives. Intron positions were determined using an in-house R script by extracting the sequences located between two consecutive exons.

### RNA extraction and RNA sequencing

For RNA sequencing, *P. bursaria* green (symbiotic) and white (aposymbiotic) cells were harvested every three hours (AM2, AM5, AM8, AM11, PM2, PM5, PM8, PM11) for RNA extraction, with a total of three replicates for each of 16 samples.

For RNA extraction, approximately 10^5^ *P. bursaria* cells were collected in early stationary phase, washed twice with 1x Dryl’s buffer, and concentrated using an 11-μm-pore-size nylon membrane. An RNeasy Mini Kit (Cat No. 74106, QIAGEN) and TRI Reagent (T9424, Sigma-Aldrich) were used to extract total RNA. An Illumina NextSeq platform was used to sequence the RNA libraries generated using an Illumina TruSeq Stranded mRNA LT Sample Prep Kit (Illumina, San Diego, CA, USA).

The adapters and low quality reads were trimmed using fastp for paired-end reads with the following parameters: cut_front_window_size=3; cut_tail_window_size=3; cut_right_window_size=4; cut_right_mean_quality=30; length_required=36 (Chen, Zhou et al. 2018). Then, the trimmed reads were aligned to the *P. bursaria* genome (Cheng, Liu et al. 2020) using STAR 2.7.11a to create BAM files with criteria: twopassMode=Basic; overSJfilterOverhangMin=7 5 5 5; outSAMstrandField=intronMotif; alignIntronMin=10; alignIntronMax=10000; alignEndsType=EndToEnd; outSAMmapqUnique=3; outSAMtype=BAM SortedByCoordinate (Dobin, Davis et al. 2013).

### Gene expression analysis

Salmon 0.13.1 was used to calculate raw gene read counts for each sample using quality-controlled FASTQ files, option gcBias, and numBoostraps=200 (Patro, Duggal et al. 2017). The tximport software was used to import raw gene counts from Salmon into R, which were then utilized in DESeq2 to normalize gene expression (Love, Huber et al. 2014, Soneson, Love et al. 2015). The differentially expressed genes (DEGs) were then analyzed using DESeq2 by computing the log2 fold-change (Log2FC) of genes between conditions and the false discovery rate (FDR) of the difference in expression level significance (Love, Huber et al. 2014). For DESeq2 to compute, the number of normalized reads for each gene must be > 10 in at least three replicates. If Log2FC ≥ 1 and FDR ≤ 0.05, then a given gene was defined as differentially expressed between two groups under consideration.

### Analyses of alternative splicing events, intron retention rate (PSI) & differential spliced intron (DSI) clustering

rMATs (replicate multivariate analysis of transcript splicing) was used to quantify alternative splicing (AS) events, including exon skipping, intron retention, mutually exclusive exons, alternative 3’ splice sites, and alternative 5’ splice sites (Shen, Park et al. 2014). Aligned BAM files from STAR were used as input for rMATs with options novelSS, allow-clipping, and mil (minimum intron size)=10 to identify the AS events between symbiotic and aposymbiotic cells. We filtered differential splicing efficiency according to |ΔPSI| ≥ 0.1, Junction Read Count (inclusion read + exclusion read) > 10 and FDR < 0.05 to define differentially spliced introns (DSI) between groups.

The individual intron retention rate was determined using SQUID (https://github.com/Xinglab/SQUID). PSI was calculated using the formula: total inclusion reads divided by total number of junction reads (Supplementary Figure 1C) (Cheng, Liu et al. 2020).

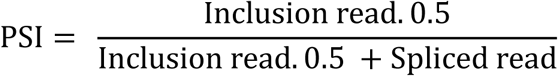

The PSI of introns was only included in our analysis if the number of Junction Read Counts was 10 or greater, and the standard deviation for PSI for three replicates was less than 0.1 in all of the replicates.

The PSI of each alternatively spliced intron between symbiotic and aposymbiotic cells was compared in a pairwise fashion at each time-point using SQUID. Introns displaying statistical significance at |ΔPSI| > 0.1 and FDR < 0.05 between symbiotic and aposymbiotic cells were selected for further investigation.

### GO term enrichment analysis

GO IDs linked to each gene ID of *P. bursaria* were identified using InterProscan 5.72-103.0, which assigned each gene to GO terms based on its protein domains (Jones, Binns et al. 2014). The clusterprofiler R tool was used for GO enrichment analysis and visualization (Yu, Wang et al. 2012). Genes with a FDR < 0.05 were considered significantly enriched in the target gene set compared to the background gene set. The enrichment score was calculated based on the ratio of genes in the target gene set divided by the ratio of genes in the background gene set.

### Spliceosome comparison using reciprocal best BLAST hit and snRNA discovery in *P. bursaria*

To compare spliceosomal components between species, data on spliceosomal proteins in *Homo sapiens* was downloaded from the Spliceosome Database as a reference (Cvitkovic and Jurica 2013). The protein fasta files of other model species (*Mus musculus*, *Danio rerio, Drosophila melanogaster*, *Caenorhabditis elegans*, *Saccharomyces cerevisiae, Arabidopsis thaliana*) were downloaded from the ENSEMBL database (Howe, Achuthan et al. 2021). *P. tetraurelia* protein fasta files were downloaded from ParameciumDB (Arnaiz, Van Dijk et al. 2017), and the protein fasta file of *Tetrahymena thermophila* was obtained from TGD Wiki (Stover, Punia et al. 2012). Homologs of human spliceosomal proteins were then identified by performing reciprocal BLAST against the protein fasta files of each species (Park, Hannenhalli et al. 2014). To improve specificity and sensitivity, reciprocal best BLAST hits (RBBHs) were allocated using both blastp and mmseq easy-rbh.

The UsnRNA multiple sequence alignment in Stockholm format was retrieved from Rfam database 14.10 (Griffiths-Jones, Bateman et al. 2003). The INFERNAL program v1.1.5 was used to create a covariance model, to calibrate the model, and to search the *P. bursaria* genome for UsnRNA sequences employing the cmbuild, cmcalibrate, and cmsearch features, respectively (Nawrocki and Eddy 2013). Significant hits against UsnRNA sequences discovered by INFERNAL were extracted using BEDTools intersect (Quinlan and Hall 2010). The sequence of each UsnRNA candidate was then sought in R2DT software, together with the corresponding template, to obtain secondary structures in DBN format (Sweeney, Hoksza et al. 2021). The structure was then annotated in RNAcanvas (Johnson and Simon 2023).

### Linear regression analysis of the relationship between PSI and splicing-related gene expression in *P. bursaria*

To elucidate the potential regulatory effect of splicing-related DEGs on identified DSIs, we performed a linear model:

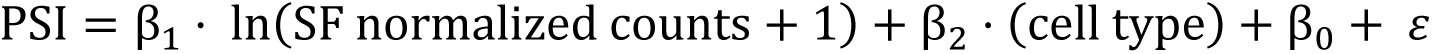

in which PSI values and ln(SF normalized counts + 1) were standardized using z-transformation across 48 replicates from 16 samples. The significance of the regression coefficients 𝛽_1_ and 𝛽_2_were assessed with FDR correction at a threshold of < 0.1. Since DSI values and splicing-related DEGs were identified from comparisons between different endosymbiotic states (symbiotic and aposymbiotic cells), we accounted for a potential cell type confounding effect by setting 𝛽_1_ to 0 when 𝛽_2_was found to be significant in the linear model. We interpreted a significant non-zero outcome for 𝛽_1_as being indicative of a potential regulatory effect of splicing-related gene expression on differential splicing.

### Intron orthologs between ciliate species and intron age assignment

The process of identifying intron orthologs was adapted from a method described previously (Olthof, Schwoerer et al. 2024). Initially, protein homologs were retrieved using mmseqs2 easy-rbh to obtain reciprocal best-hit (RBH) proteins between species (Steinegger and Soding 2017). Next, the amino acid sequences of exon-exon junctions, comprising the 10 amino acids on either side of the junction, were extracted to define intron positions. These intron position sequences from protein homologs were then subjected to pairwise alignment using the pairwiseAlignment function in the Biostrings R package, using the BLOSUM-62 scoring system. Intron alignments were considered RBHs if their aligned regions exhibited at least 40% sequence identity and achieved the highest alignment score in both forward and reverse directions.

For introns found in orthologous genes shared at least between three out of four examined species, we determined their evolutionary age using phylostratigraphy and the maximum parsimony method (Domazet-Loso, Brajkovic et al. 2007, Kannan and Wheeler 2012). For this analysis, the youngest intron group, assigned an age of 1, is specific to the reference species, whereas the oldest intron group consists of introns originating from the first node in the phylogenetic tree examined.

## Data availability

The transcriptomic data generated for and analyzed in the current study are available from NCBI under the accession number BioProject PRJNA1279681 (https://dataview.ncbi.nlm.nih.gov/object/PRJNA1279681?reviewer=i5u6euocb0lp8fe6nc9e69ht <u>6g</u>). Codes used for the analyses in this study have been deposited at https://doi.org/10.5281/zenodo.16736398

## Results

### The extremely short introns in *P. bursaria* enhance gene expression

To characterize the features of *P. bursaria* introns, we annotated 39,715 introns in the functional gene set from the DK2 genome annotation file (Supplementary Table 1, see Materials and Methods for details). More than 80% of *P. bursaria* genes contain at least one intron, with an average of 2.7 introns per gene (Supplementary Figure 1A). Moreover, we discovered a unique distribution of introns in *P. bursaria* transcripts, being enriched at both the 5’ and 3’ gene ends (Figure 1A). A bias of introns towards the 5’ end of transcripts has also been reported for other model species, such as the budding yeast, fruit fly and mouse (Sakurai, Fujimori et al. 2002), implying a general feature of eukaryotic introns. A majority of *P. bursaria* introns are less than 40 nucleotides (nt) in length, with a median length of 24 nt, and minimum and maximum lengths of 15 and 100 nt, respectively (Supplementary Figure 1B). The distribution of intron lengths is very similar to that of *Paramecium tetraurelia*, which displays a median intron length of 25 nt (Arnaiz, Van Dijk et al. 2017). In Supplementary Figure 1C, we present a sequence logo plot showing that most of the annotated introns in *P. bursaria* are canonical introns with conserved GT-AG 5’ and 3’ splice boundaries, indicating that an exclusively U2-dependent splicing mechanism operates. Although the GC content of *P. bursaria* exons is ∼30.6%, the GC content of its introns is substantially lower (average = 17.7%) (Supplementary Figure 1D).

**Figure 1.**
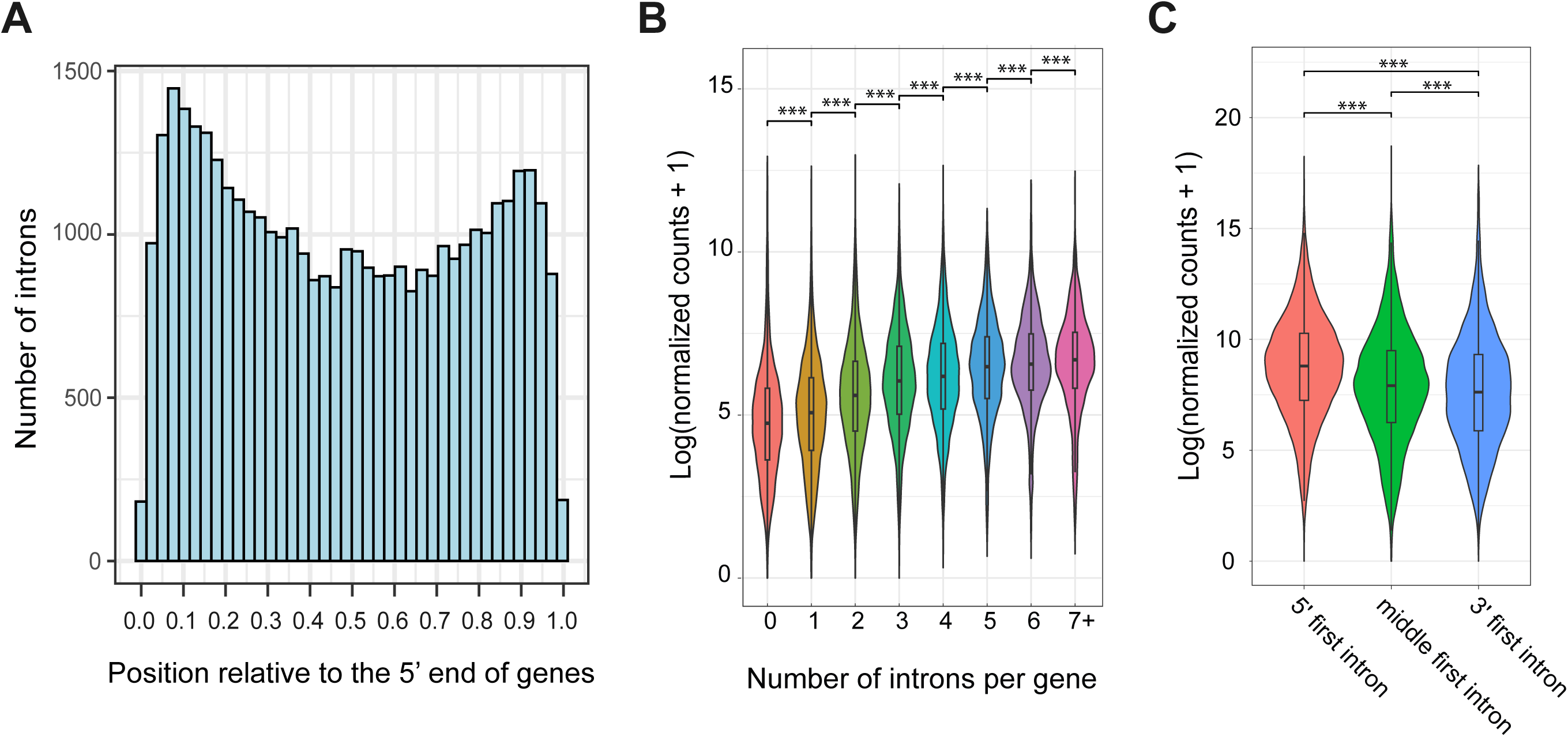
Intron position and intron-mediated enhancement of gene expression in *P. bursaria.* A. The distribution of introns is enriched at both the 5’ and 3’ ends of genes. Distribution of introns relative to the 5’ end of protein-coding regions. Intron positions were determined by dividing the distance from the intron to the first codon by the total gene length. B. Gene expression levels are positively correlated with the number of introns per gene. C. Genes having the first intron at their 5’ end display significantly higher expression than the other two groups. Genes were grouped according to the position of their first intron. ***, p-value < 0.001, Mann-Whitney U test.

To compare how splicing is physiologically regulated between symbiotic and aposymbiotic cells, we performed transcriptomics analysis on samples collected from both cell types at eight time-points (8 AM, 11 AM, 2 PM, 5 PM, 8 PM, 11 PM, 2 AM, and 5 AM) through a light-dark cycle. We used the resulting data to explore the characteristics of small intron splicing in *P. bursaria*. In eukaryotes, intron splicing plays a crucial role in regulating gene expression through the co-transcriptional splicing mechanism (Shaul 2017). We investigated if the small introns in *P. bursaria* exert a similar function. Transcriptomic data from all 16 samples were pooled together and analyzed. We observed that genes containing introns exhibited significantly higher expression levels than those without introns (Wilcoxon rank-sum test, p < 0.001). Moreover, gene expression level was positively correlated with intron number (Figure 1B).

This positive correlation between intron numbers and gene expression levels in *P. bursaria* implies intron-mediated gene expression enhancement. U1 snRNPs can directly recruit transcription factors, such as TFIIH, TFIIB, and TFIID, to a gene promoter when the intron 5’ splice site is located near the promoter (Das, Yu et al. 2007, Damgaard, Kahns et al. 2008). Since we detected a bias in intron positions toward the 5’ end of transcripts (Figure 1A), we investigated if the presence of introns at this position is associated with enhanced gene expression in *P. bursaria*. To do so, we grouped and compared *P. bursaria* genes based on the relative position of the first intron: the 5’ end group (i.e., the first intron is located within 0-25% of the total length), the middle group (within 25-75% of total length), or the 3’ end group (within 75-100% of total length). Indeed, the 5’ end group displayed significantly higher expression than the other two groups (Figure 1C).

### Conservation of the U2-dependent spliceosome in *P. bursaria*

Compared to other eukaryotes having well-characterized splicing mechanisms (Zhu, He et al. 2010, Yang, Beutler et al. 2022), intron size in *P. bursaria* is relatively small (Supplementary Figure 1B). Spliceosomes are huge protein complexes containing many snRNAs and proteins. The human spliceosome A complex is ∼205×195×150 Å in size and covers 79-125 nt of a stretched RNA, based on its cryogenic electron microscopy (cryo-EM) structure (Behzadnia, Golas et al. 2007). The *P. bursaria* spliceosome likely requires extensive reorganization to function on small introns. To investigate this possibility, we downloaded 1,005 human splicing-related proteins from SpliceosomeDB (Cvitkovic and Jurica 2013). We identified orthologs across three representative ciliate species (*P. bursaria*, *P. tetraurelia*, and *T. thermophila*), along with other model organisms, including *M. musculus*, *D. rerio*, *D. melanogaster*, *C. elegans*, *S. cerevisiae*, and *A. thaliana* (Supplementary Table 2) (Steinegger and Soding 2017). Among all of the species we analyzed, 275 out of 1,005 spliceosomal proteins were shared, representing core spliceosome components (Figure 2A).

**Figure 2.**
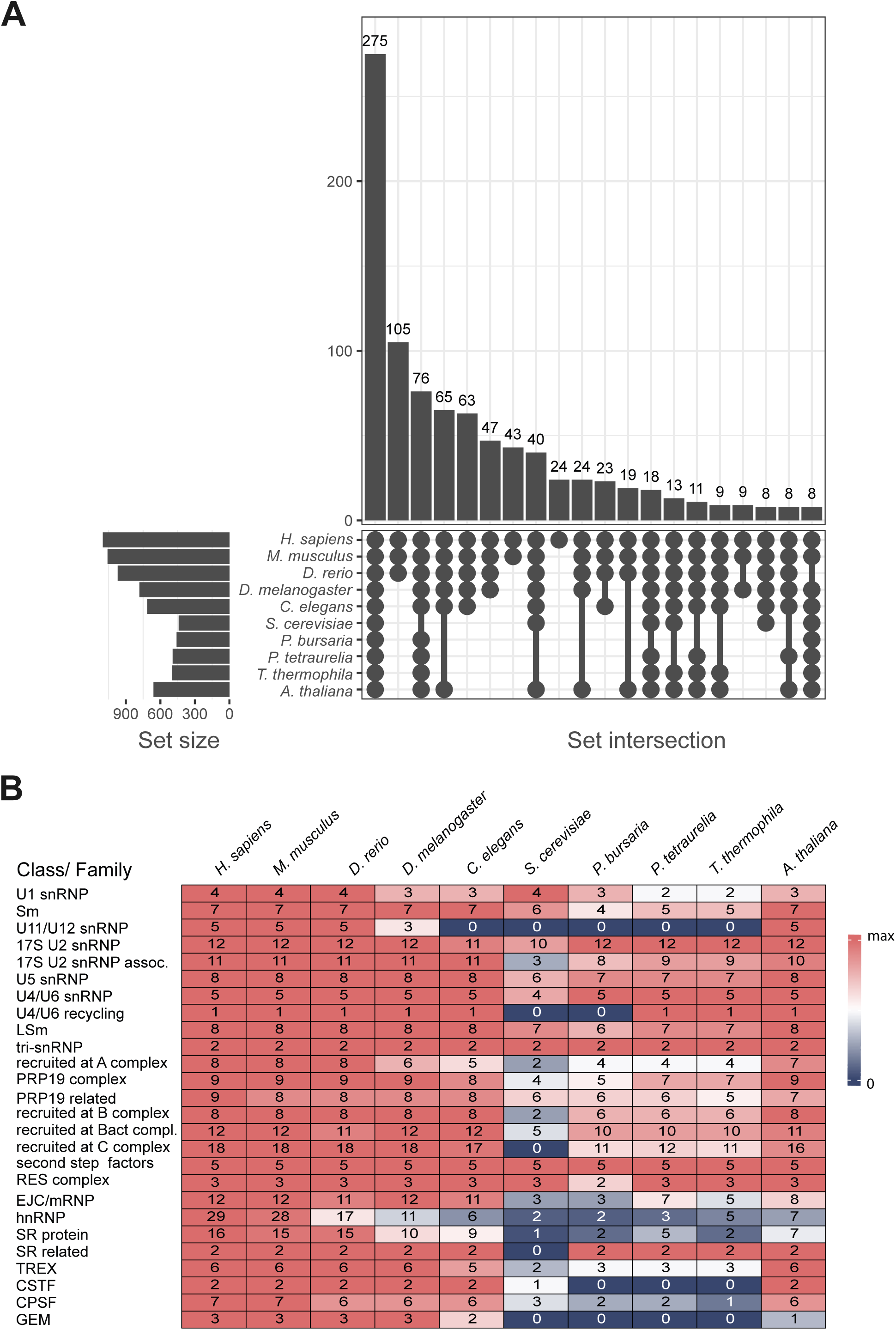
Conserved splicing-related proteins in model organisms and ciliate species. A. An UpSet plot displaying the numbers of splicing-related proteins shared among 10 species. Set size represents the number of splicing-related proteins in each species, with numbers above the bars indicating the count of proteins in each shared group. B. Number of splicing-related proteins in each protein class/family across 10 species. Colors indicate protein counts relative to the median: red (above median), white (equal to median), and blue (below median).

Forty human splicing-related proteins were not identified in the ciliate species, including a U1 snRNP (LUC7L), a U5 snRNP (CD2BP2), and an LSm protein (LSM1) (Figure 2A, Supplementary Table 3). In human cells, LUC7L enhances splicing by recognizing a strong consensus sequence downstream of the 5’ splice site (within the intron), whereas its paralog LUC7L3 promotes splicing by interacting with an upstream consensus sequence located within the exon (Daniels, Hershberger et al. 2021, Kenny, McGurk et al. 2025). Loss of LUC7L and retention of LUC7L3 in *P. bursaria* corresponds to it possessing a conserved upstream consensus (positions-2 and-1 in the exon) and lacking a downstream consensus (positions +3 to +6 in the intron) to the 5’ splice site (Supplementary Figure 1C), implying a streamlined mechanism of 5’ splice site recognition by the U1 snRNP, likely a consequence of the reduced splicing machinery possessed by ciliates.

Interestingly, we observed a complete absence of the U11/U12 snRNP, CSTF, and GEM complexes among the ciliates we considered (Figure 2B). Additionally, we uncovered a reduction in the number of proteins associated with regulatory classes, including SR proteins, hnRNPs, and CPSF (Figure 2B). Notably, although the numbers of splicing-related proteins in ciliates and *S. cerevisiae* are relatively similar, yeast and ciliate species possess different sets of unique proteins (Figure 2A and Supplementary Table 3), likely reflecting different trajectories of spliceosome reorganization, one pertaining to the reduction in intron number displayed by *S. cerevisiae* and another for the reduction in intron size of ciliates.

Overall, the numbers of U2-dependent core spliceosome proteins in *P. bursaria*, including U1, U2, U4, U5, U6, Sm, and Lsm proteins, are conserved. To further investigate the snRNA repertoire in *P. bursaria*, we analyzed conserved secondary structures of UsnRNAs from the Rfam database (Griffiths-Jones, Bateman et al. 2003). We detected U1, U2, U4, U5, and U6 snRNAs with conserved secondary structures and key functional motifs, but not the U11 or U12 snRNAs (Supplementary Figure 2). Together, our data indicate that a modified U2-dependent splicing mechanism operates in *P. bursaria* to splice small introns.

### Intron retention and alternative 3’ splice sites are the most abundant alternative splicing events in *P. bursaria*

Since alternative splicing influences both protein isoform diversity and gene expression levels, it may play a crucial role in adaptive regulation of the endosymbiotic process. To explore this possibility, we analyzed how five major types of alternative splicing events are regulated across eight time-points between symbiotic and aposymbiotic *P. bursaria* cells: intron retention (RI), exon skipping (SE), alternative 3’ splice site (A3SS), alternative 5’ splice site (A5SS), and mutually exclusive exon (MXE) (Supplementary Figure 3). Our results show that intron retention and alternative 3’ splice site events were the most frequently regulated between symbiotic and aposymbiotic cells (Figure 3A). We then filtered for significant A3SS and RI events (FDR < 0.05) using junction read counts (total junction reads ≥ 10) and Percent Splice In (PSI) changes (|ΔPSI| ≥ 0.1), which uncovered 512 significant A3SS events (Figure 3B) and 992 significant RI events (Figure 3C).

**Figure 3.**
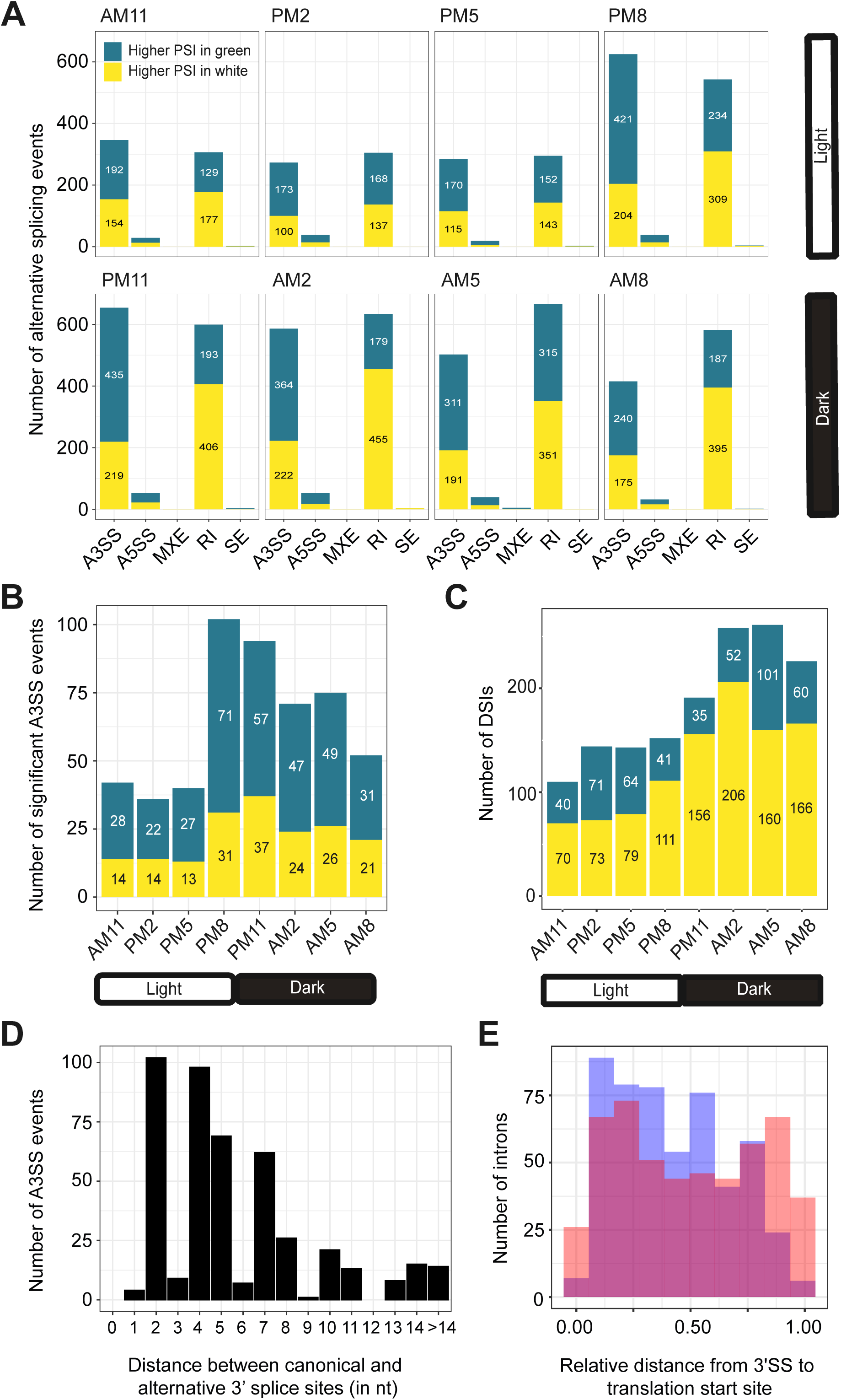
Alternative splicing events between symbiotic and aposymbiotic *P. bursaria* cells. A. Intron retention and alternative 3’ splice site events are the most frequently regulated between symbiotic and aposymbiotic cells. Numbers of significant alternative splicing events (FDR < 0.05) between symbiotic and aposymbiotic cells at eight time-points through a single 24-hour light-dark cycle. Green indicates higher PSI in symbiotic cells and yellow indicates higher PSI in aposymbiotic cells. The alternative splicing events are A3SS: alternative 3’ splice site, A5SS: alternative 5’ splice site, MXE: mutually exclusive exon, RI: intron retention, and SE: exon skipping. B. Number of significant alternative 3’ splice site (A3SS) events between symbiotic and aposymbiotic cells. C. Number of differentially spliced introns (DSIs) between symbiotic and aposymbiotic cells. D. Distance (in nucleotides) between the alternative 3’ splice site to the canonical 3’ splice site for significant A3SS events between symbiotic and aposymbiotic cells. E. Introns displaying A3SSs are significantly enriched in the 5’ portion of genes (Kolmogorov-Smirnov test, p-value = 7.599e-07). Blue indicates introns with a A3SS and orange represents the same number of introns randomly selected from among all introns.

Interestingly, significant A3SS and RI events proved to be more frequent at night (PM11, AM2, AM5, AM8) than during the day (AM11, PM2, PM5, PM8) (Figure 3B and 3C). Moreover, during night time-points, a higher ratio of A3SS events presented increased PSI in symbiotic cells relative to aposymbiotic cells (Figure 3B). In contrast, RI events exhibited a higher proportion of increased PSI in aposymbiotic cells at night (Figure 3C). Since *P. bursaria* cells exhibit rhythmic physiological changes during the day-night cycle (Miwa 2009), our results indicate that alternative splicing may be involved in these changes.

To assess the functional impact of the alternative splicing events, we investigated the impact of A3SS events on the translation reading frame by calculating the distance between canonical 3’SS and alternative 3’SS. Unexpectedly, we found that the majority of distances were not multiples of nucleotide triplets (n=479 out of 512 events), which would cause a frameshift in open reading frames and potentially trigger mRNA degradation via the NMD pathway (Figure 3D).

This outcome differs from the 3’ wobble splicing observed in mammalian cells (e.g., NAGNAG motifs), which often preserves functional reading frames from both 3’SS choices (Hiller, Huse et al. 2004, Akerman and Mandel-Gutfreund 2006). Furthermore, we observed that A3SS events occur more often in the introns located in the 5’ portion of genes (Figure 3E). In addition, exon junction presence is significantly more common downstream of A3SS event introns compared to random introns (odds ratio=1.97, Fisher’s exact test p-value=1.101e-06). Taken together, these results indicate that A3SS selection in *P. bursaria* predominantly serves as a mechanism to abrogate protein production.

### Intron retention rates are associated with GC content, intron length and gene expression levels

Next, we investigated the intrinsic factors influencing intron retention rate in *P. bursaria*. After filtering out the introns with low junction read counts (i.e., read count ≤ 10), 66% of total introns (n=26,123 out of 39,715) presented sufficient counts to calculate splicing efficiency in at least one sample. More than 87% of these introns exhibited intron retention rates (presented as PSI) lower than 0.1 for all samples, indicating that most introns in *P. bursaria* are spliced efficiently (Supplementary Figure 4A and Table 4).

We detected a strong positive correlation between the GC content of introns and intron retention rates (Figure 4A). Although a similar trend has been reported previously for *P. tetraurelia* (Gnan, Matelot et al. 2022), the effect of GC content on intron retention appears to be much stronger in *P. bursaria*. In addition to intron GC content, we found that intron length also contributes to splicing efficiency. Introns larger than 26 nt or smaller than 23 nt were spliced less efficiently than introns of 23-26 nt, representing the middle 50% of all introns (Figure 4B).

**Figure 4.**
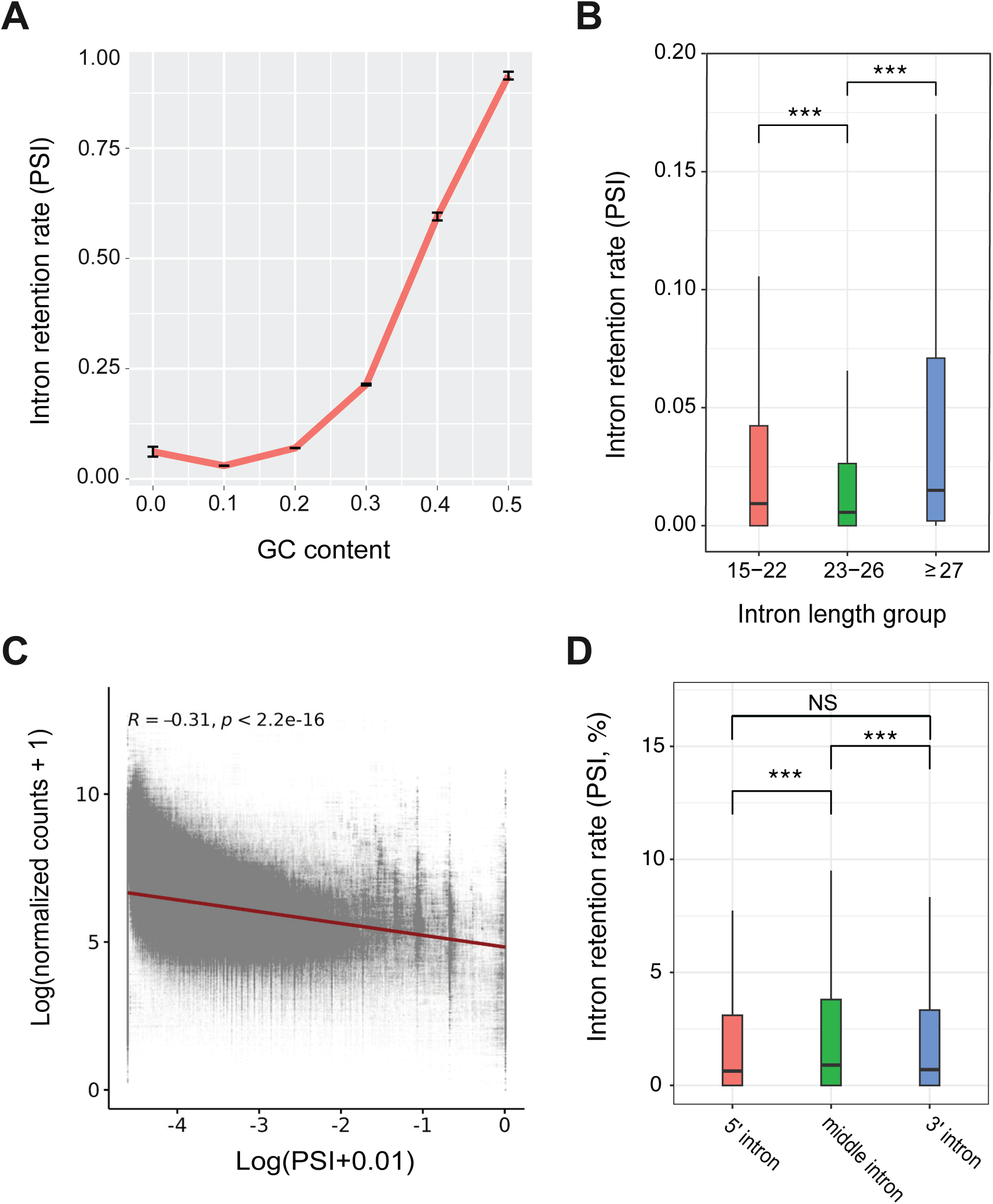
The intron retention rate correlates with intron sequence features and gene expression. A. The intron retention rate is positively correlated with the GC content of introns. GC content values have been rounded to the nearest group. Error bars indicate standard error of the mean. B. Intron retention rates across different intron length groups. The introns were divided into three groups, 15-22 nt, 23-26 nt, and ≥ 27 nt, representing approximately the first, middle two, and last quartiles of all introns. ***, p-value < 0.001, Wilcoxon rank-sum test. C. Gene expression levels are negatively correlated with intron retention rates, based on a Pearson correlation analysis. D. 5’ end and 3’ end introns are spliced more efficiently than middle introns. Introns were classified as 3’ introns (relative position ≥ 0.75), 5’ introns (relative position ≤ 0.25), or middle introns (0.25 < relative position < 0.75). ***, p-value < 0.001; NS, p-value > 0.05, Mann-Whitney U test.

As shown in Figure 1, genes with a higher intron count and the first intron positioned near the 5’ end of genes tend to display elevated expression levels. In addition, we observed a negative correlation between expression level and the intron retention rate (Figure 4C), indicating that efficient splicing contributes to transcript abundance. Moreover, introns at the 5’ and 3’ ends are spliced more efficiently than introns in the middle of genes (Figure 4D), consistent with their positional enrichment and potential roles in regulating transcript maturation (Figure 1A and 1C). These results highlight that intron splicing, especially for introns located at the 5’ end of genes, is associated with enhanced gene expression in *P. bursaria*, as also reported previously for mammalian cells (Nott, Meislin et al. 2003, Park, Hannenhalli et al. 2014).

### Symbiotic and aposymbiotic cells display two distinct intron splicing patterns associated with differential expression of splicing factors

To investigate the intron splicing pattern further, we compiled all differentially spliced introns (DSIs) (|ΔPSI| ≥ 0.1 and FDR < 0.05) from all time-points (Figure 3C), resulting in a total of 992 DSIs in 883 genes (Supplementary Table 5). A principle component analysis (PCA) was then performed to cluster the samples (16 samples, each with three replicates) based on the PSI values of these DSIs. The PSI profiles of the DSIs effectively distinguished symbiotic from aposymbiotic cells (Figure 5A), indicating that endosymbiosis influences the splicing pattern. Additionally, symbiotic cells formed two sub-clusters along PC2, one corresponding to nighttime, and the other including cells from the daytime and the first nighttime time-point (PM11) (Figure 5A), possibly linked to photosynthesis by the endosymbiotic algae.

**Figure 5.**
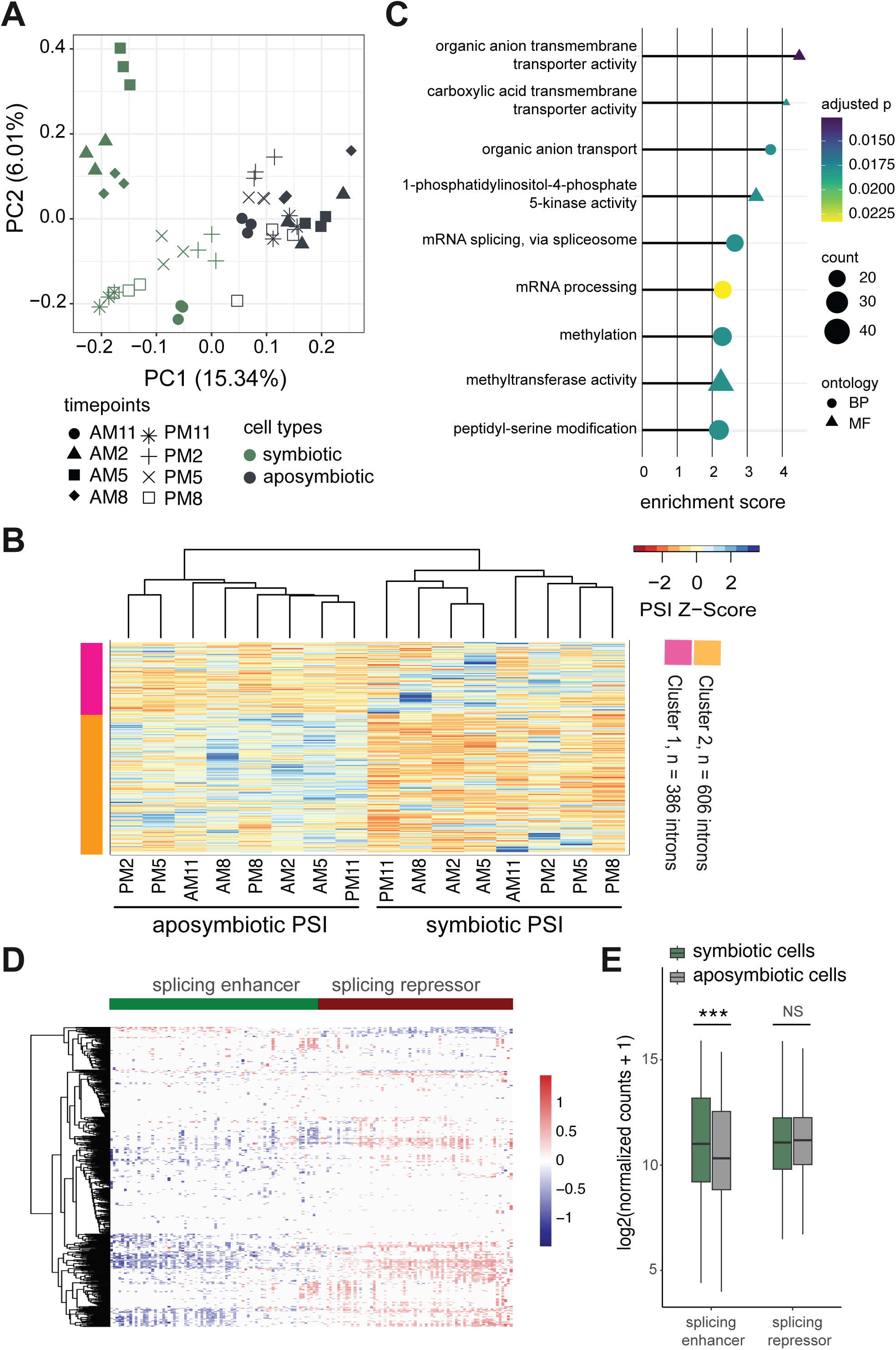
Patterns of intron splicing between symbiotic and aposymbiotic cells. A. Symbiotic and aposymbiotic samples are clearly separated by principal component analysis on the intron retention rates of 992 DSIs. Each time-point sample comprised three biological replicates. B. k-means clustering separates the identified DSIs into two clusters displaying distinct patterns of intron retention (PSI). The heatmap shows the PSI values for 16 samples in two DSI clusters. Color is scaled according to PSI z-score. Cluster 1 (pink), n = 386 introns; cluster 2 (orange), n = 606 introns. C. GO term enrichment for genes containing DSIs. D. Association of DSIs with the expression levels of splicing-related genes. Heatmap shows the coefficients (β_1_) of splicing-related genes from the linear model of DSI PSI values, based on 992 DSIs (rows) and 147 differentially expressed splicing-related genes (columns). These splicing-related genes can be generally classified into two distinct groups, i.e., splicing enhancers and repressors. E. Splicing-enhancing factors are significantly upregulated in symbiotic cells. Gene expression levels of splicing enhancers and repressors in symbiotic and aposymbiotic cells are shown. ***, p-value < 0.001; NS, p-value > 0.05, Wilcoxon rank-sum test.

Then, we performed k-means clustering to separate the identified DSIs into two clusters having distinct intron PSI patterns (Figure 5B, Supplementary Table 5). Cluster 1 consists of DSIs with higher PSI in symbiotic cells (n=386 introns), whereas cluster 2 (n=606 introns) includes DSIs with higher PSI in aposymbiotic cells (Figure 5B). Gene ontology (GO) term enrichment analysis of genes containing DSIs revealed an overrepresentation of mRNA splicing and transport-related terms, including organic anion transmembrane transporter activity, organic anion transport, and carboxylic acid transmembrane transport (Figure 5C and Supplementary Table 6). Previous studies have shown that endosymbiotic algae are maintained in a membranous compartment (i.e., the perialgal vacuole) within *P. bursaria* host cells, and that the host and endosymbionts continuously exchange several organic compounds to establish a stable mutualistic relationship (Fujishima and Kodama 2012). Thus, our findings indicate that intron splicing may play a pivotal role in regulating these transmembrane exchanges. Moreover, since intron retention rates in spliceosome genes appear to be regulated, there is potentially autoregulation of splicing-related genes in the context of endosymbiosis, which may in turn influence the retention levels of other introns.

Certain splicing factors can regulate the introns of genes involved in specific cellular pathways. For instance, HNRNPK and SRSF1 were shown previously to control intron retention during B cell development (Ullrich and Guigo 2020). Since we found that intron splicing efficiency is regulated in endosymbiosis, potential splicing factors may influence splicing of DSIs during this process. To explore this supposition, we analyzed 6,949 differentially expressed genes (DEGs) between symbiotic and aposymbiotic cells across all time-points and identified 147 splicing-related genes (see Materials and Methods for details). Next, we used a linear regression model to elucidate the potential regulatory effect of the 147 splicing-related DEGs on 992 DSIs.

Since we identified both differentially expressed splicing factors and DSIs in our comparisons of symbiotic and aposymbiotic cells, the status of endosymbiosis may confound any associations. To account for this issue, we incorporated endosymbiotic status as a covariate in the model (DSI ∼ splicing-related DEG + endosymbiotic status) and retained only DSIs whose coefficients were significant with splicing-related DEGs but not for endosymbiotic status (see Materials and Methods for details). Applying this filter yielded 477 introns, which likely represent direct regulatory targets of these splicing factors (Figure 5D and Supplementary Table 7). We could classify the 147 splicing-related genes into two groups based on their mean correlation coefficient with intron retention rates (PSIs): splicing-enhancing factors, whose mean coefficient with the PSI of DSIs is negative, and splicing-repressing factors, whose mean coefficient is positive with PSI (Figure 5D). Notably, splicing-enhancing factors were significantly upregulated in symbiotic cells (Figure 5E), implying that symbiotic cells regulate these splicing-related factors to fine-tune the splicing and expression of intron-containing genes.

### Intron evolution in ciliate species reveals higher splicing efficiency and lower intron GC content in aged introns

We observed that some intron characteristics, such as GC content and the intron distribution within genes, are closely associated with splicing efficiency and gene expression regulation (Figures 1 and 3). Moreover, symbiotic cells may adjust the expression of specific splicing factors to regulate splicing efficiency and mRNA levels (Figure 5). Given the functional significance of introns, we explored the evolutionary patterns of introns within Genus *Paramecium*.

For this analysis, we selected three *Paramecium* species with well-annotated genomes, i.e., *P. bursaria, P. tetraurelia,* and *P. caudatum*, with *T. thermophila* acting as an outgroup. We identified 8,886 orthologs shared across these species. We then identified the best-matching introns from paired orthologous genes between *P. bursaria* and the other species (see Materials and Methods for details).

To classify intron origins, we applied phylostratigraphy and maximum parsimony, assigning introns to different nodes on the phylogenetic tree. We observed that 10.1% of introns in orthologous genes belong to the oldest group (n=2,680 introns, age group=3), with 43.3% of introns being specific to *Paramecium* (n=11,475, age group=2). The remaining 46.6% of introns are unique to *P. bursaria* (n=12,336, age group=1), highlighting dynamic evolution of introns in this lineage (Figure 6A).

**Figure 6.**
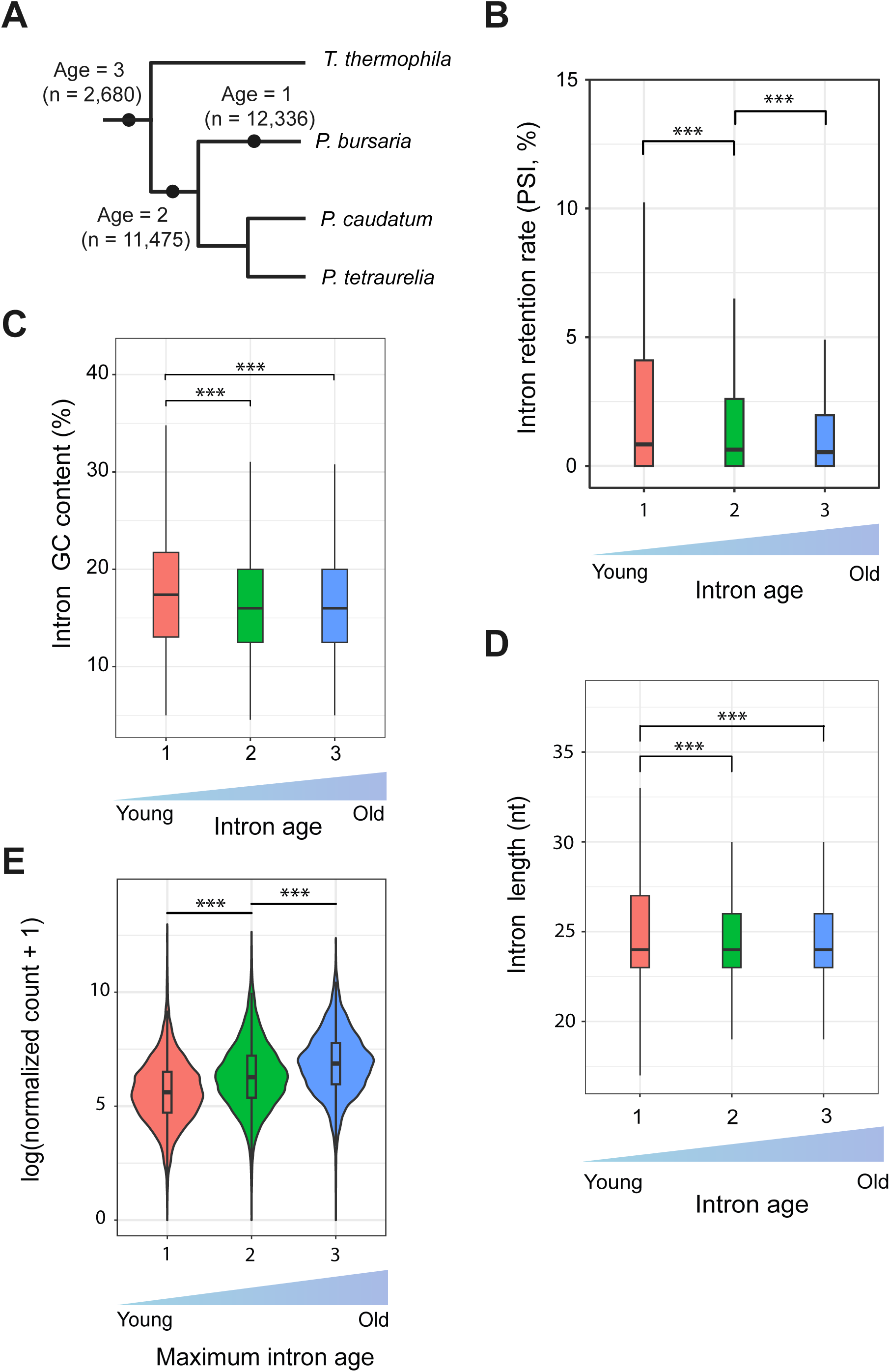
Short intron evolution in *P. bursaria*. A. A diagram showing the phylogenetic relationships between different *Paramecium* and *Tetrahymena* species. Introns in *P. bursaria* were assigned to different age groups based on their conservation between *P. bursaria* and the other species. The Age 1 group represents the youngest introns, whereas the Age 3 group represents the oldest introns. B. Young introns exhibit greater intron retention than old introns. Boxplots show the intron retention rate for each intron age group. ***, p-value < 0.001, Mann-Whitney U test. C. Young introns display a higher GC content than old introns. Boxplots show the intron GC content for each intron age group. ***, p-value < 0.001, Mann-Whitney U test. D. Young introns present a wider intron size distribution than old introns. Boxplots show intron length (in basepairs) for each intron age group. ***, p-value < 0.001, Kolmogorov-Smirnov test. E. Genes containing old introns are more highly expressed than those solely possessing young introns. Boxplots show expression level for each gene group based on the maximum intron age in the genes. ***, p-value < 0.001, Mann-Whitney U test.

Interestingly, older introns (age groups 2 and 3) exhibit better splicing efficiency than newer introns (age group 1) (Figure 6B). In *P. bursaria*, high-GC introns generally have higher PSI values compared to low-GC introns. This scenario aligns with our observation that new introns in *P. bursaria* have higher GC contents than older ones (Figure 6C). We also observed a trend whereby younger introns present a wider range of length distributions compared to introns in the older groups (Figure 6D). Consistently, we detected that introns longer than 27 nt or shorter than 23 nt are spliced less efficiently compared to those within the 23–26 nt range (Figure 4B), indicating that the intron length distribution in *P. bursaria* has evolved over time toward an optimal range. Given the positive association between intron presence, intron number, and gene expression, we assessed gene expression in relation to intron age, reflecting when introns were acquired in the evolutionary tree. We categorized the 8,886 orthologous genes into three groups based on the age of their oldest intron: group 1 comprised genes with only young introns (n=1,466); group 2 included genes with a maximum intron age of 2 (n=4,863); and genes with a maximum intron age of 3 were assigned to group 3 (n=1,656). Notably, genes hosting older introns exhibited significantly higher gene expression levels compared to those with only young introns (Figure 6E). This finding further supports a role for introns in enhancing gene expression.

To determine if the evolutionary patterns we observed in *Paramecium* introns are also present in other ciliates, we extended our analysis to *Tetrahymena*. We selected four representative species from Genus *Tetrahymena*, including *T. thermophila, T. malaccensis, T. eliotti* and *T. borealis*. Using the same approach as performed for *Paramecium*, we classified introns in *T. thermophila* that had gene orthologs in at least three *Tetrahymena* species into four intron age groups (Supplementary Figure 5A).

Our results revealed that the majority (68.9%) of introns in *T. thermophila* belonged to the oldest group (n=42,030 of a total of 60,971 introns). In contrast, species-specific introns (age group 1) in *T. thermophila* made up only 9.3% of the total introns in shared orthologous genes (n=5,678 introns) (Supplementary Figure 5A). We found that younger introns also exhibited a higher retention rate, a pattern consistent with our result for *P. bursaria* introns (Supplementary Figure 5B). Similarly, young introns in *T. thermophila* had a higher GC content and a wider length range compared to older introns (Supplementary Figures 5C and 5D). When we categorized genes based on their oldest intron age in *T. thermophila*, we observed a significant trend whereby genes containing older introns exhibited higher expression levels compared to those with only younger introns (Supplementary Figure 5E). Taken together, our analysis of intron features across different evolutionary age groups reveals a consistent pattern of intron optimization, characterized by improved splicing efficiency and enhanced gene expression, during evolution of both the *Paramecium* and *Tetrahymena* lineages.

## Discussion

### Alternative splicing as a mode for gene regulation in endosymbiosis

Host cells integrating symbionts likely experience drastic changes in gene expression. A previous study on *P. bursaria* uncovered nearly 7,000 DEGs between symbiotic and aposymbiotic cells (Yuuki Kodama 2014), which is consistent with our results presented herein. Another study indicated that upregulation of glutamine biosynthesis is correlated with symbiont abundance (He, Wang et al. 2019), supporting that gene regulation plays a crucial role in endosymbiosis. Similar phenomena have been observed in other examples of algal secondary endosymbiosis. For example, genes involved in glutamine synthesis and phosphate transporters are also upregulated in *Hydra viridissima* hosting algal endosymbionts (Hamada, Schroder et al. 2018). In addition to transcriptional regulation, post-transcriptional regulation, such as alternative splicing, can provide the host with more flexibility in adjusting the endosymbiotic relationship, especially when the endosymbiotic algae undergo physiological changes relating to photosynthesis during the day/night cycle. A recent study on *P. bursaria* found that 6mA DNA methylation facilitates intron retention and may serve as a reverse marker to distinguish endosymbiosis-related genes (Pan, Ye et al. 2023), providing a potential link between splicing regulation and endosymbiosis. Here, our study shows that 992 introns are differentially spliced between symbiotic and aposymbiotic *P. bursaria* cells (Figure 5), further supporting that splicing regulation is involved in endosymbiosis.

### Splicing factors specifically regulate DSIs between symbiotic and aposymbiotic cells

Alternative splicing is regulated by many factors, such as co-transcriptional, chromatin structure, DNA sequence, and epigenetic modifications, as well as splicing factors (Chen and Manley 2009). In *P. tetraurelia*, nucleosome positioning affects intron splicing, with introns at the edges of nucleosomes displaying greater splicing efficiency (Gnan, Matelot et al. 2022). Moreover, 6mA DNA methylation in *P. bursaria* was shown previously to be enriched in retained introns (Pan, Ye et al. 2023). Alternative splicing factors are regulated in a tissue-specific manner in humans. For example, expression of polypyrimidine tract binding protein 1 (PTB1) is high in neuron progenitor cells, but significantly downregulated in differentiated neurons (Boutz, Stoilov et al. 2007, Keppetipola, Sharma et al. 2012). Since endosymbiosis induces many physiological changes in the host cell, including the formation of new organelles for endosymbionts, alternative splicing regulation during this process may be influenced by specific splicing factors. Our study has identified differentially expressed splicing-related genes between symbiotic and aposymbiotic cells, implying a potential role for these splicing factors in intron splicing during endosymbiosis. Furthermore, a linear regression model between splicing-related DEGs and differentially spliced introns (DSIs) across 16 samples revealed that a subset of the DSIs is strongly associated with those splicing-related DEGs. Two groups of splicing factors possibly represent “splicing enhancers” and “splicing repressors” of those DSIs in the endosymbiotic relationship (Figure 5D).

To pinpoint key splicing factors strongly associated with the PSI of DSIs, we ranked 147 splicing factor DEGs based on the number of significant coefficients across introns (Figure 5D, Supplementary Figure 4B). The top 30 splicing-related genes include several ribosomal proteins and core splicing factors, such as U1 snRNPs (SNRNP70, SNRPC), an Sm protein (SNRPD1), a second-step spliceosome factor (PRPF18), RNA helicases (DDX23, DDX46), a U5 snRNP (TXNL4A), and chromatin-related proteins (SMARCA5, SMARCA1).

The expression of several ribosomal proteins is negatively associated with the PSI of potential regulated introns, including RPL22, RPS20, and RPS11. Apart from their canonical function in protein translation, ribosomal proteins have also been shown to exert alternative splicing regulatory activity. For example, the large protein subunit L10a in nematodes and vertebrates can bind to its own pre-mRNA to switch splice site choice (Takei, Togo-Ohno et al. 2016). Although the exact regulatory functions of such ribosomal proteins remain to be studied further, it is possible that they act in alternative splicing regulation by binding to RNA or by interacting with other splicing factors.

For the top 30 genes associated with DSIs, we found that the coefficient relationships (i.e., either positively or negatively related to DSIs) of splicing factors aligned well with their known functions in splicing regulation. For instance, PRPF18 plays a crucial role in maintaining high splicing fidelity during the second splicing step, and G3BP2 interacts with PSF (polypyrimidine tract-binding protein-associated splicing factor) to enhance mRNA stability (Han, Liu et al. 2022, Roy, Gabunilas et al. 2023, Takayama, Suzuki et al. 2024). However, we also found that the coefficient relationship for some genes differed from their functions suggested by previous studies. For example, PRPF4B phosphorylates the SRSF1 protein, allowing it to work as a splicing enhancer in the fission yeast (Chen, Moore et al. 2007), yet we found that PRPF4B acts as a splicing repressor on DSIs in *P. bursaria*. Thus splicing regulation appears to operate differently in *P. bursaria*, particularly in the context of endosymbiosis. Further research is necessary to explore how the regulation of DSIs is involved in endosymbiosis.

### Intron evolution is closely linked to gene expression regulation

Although most introns are spliced efficiently in *P. bursaria*, we observed a significant difference between the splicing efficiency of old and newly acquired introns (Figure 6). We detected a similar pattern for *Tetrahymena* (Supplementary Figure 5). This outcome indicates that introns gained over time can adapt to the splicing system, potentially through sequence modifications or changes in splicing factor binding or interactions (Yeo, Van Nostrand et al. 2007, Schirman, Yakhini et al. 2021). A previous study demonstrated that intron orthologs from various species lacking U2AF1 and having a shorter distance between the branch point site (BPS) and the 3’ splice site (3’SS) are spliced more efficiently in *S. cerevisiae* (which also lacks U2AF1 and has a short BPS-to-3’SS distance) compared to intron orthologs from species that possess U2AF1 orthologs and have a longer BPS-to-3’SS distance. This finding underscores specialization of the splicing machinery of a particular species for its own intron structures (Schirman, Yakhini et al. 2021). Our analyses consistently uncovered differences between new and old intron sequences, showing that new introns in *P. bursaria* have a higher GC content and a wider length distribution than old introns (Figure 6). Intron GC content was shown previously to be negatively correlated with splicing efficiency in both *P. tetraurelia* and *S. cerevisiae* (Schirman, Yakhini et al. 2021, Gnan, Matelot et al. 2022). We report herein a similar trend in *P. bursaria* (Figure 4A). In terms of intron length, those of 23-26 nt in *P. bursaria* exhibit greater splicing efficiency than longer or shorter introns (Figure 4B). A wider length distribution of new introns could also contribute to the reduced splicing efficiency we detected.

We also observed higher expression of genes containing old introns compared to genes only having new introns (Figure 6E). Moreover, gene expression levels were positively correlated with intron number (Figure 1B). Thus, it is clear that the presence of introns is tightly associated with gene expression levels. The reduced splicing efficiency of new introns may be attributable to new introns still undergoing optimization specific to the splicing machinery. Consequently, through further sequence modifications, they may ultimately behave like old introns. Alternatively, some new introns may have specific regulatory functions, providing the cell with a strategy to fine-tune splicing efficiency (and therefore gene expression) under different conditions.

## Acknowledgments

We thank members of the Lin and Leu labs for helpful discussion and comments on the manuscript. We also thank John O’Brien for manuscript editing and the IMB Genomics Core for experimental assistance. This work was supported by Academia Sinica of Taiwan (grant no. AS-IA-110-L01 and AS-GCS-113-L03) and the National Science and Technology Council of Taiwan (NSTC 113-2326-B-001-002 and NSTC 112-2628-B-001-009). KMM was supported by an NSTC postdoctoral fellowship (NSTC 113-2811-B-001-065).

## Author contributions

CLL and JYL conceived the study. TNGN, KMM, CLL, and JYL designed analyses and interpreted results. TNGN and KMM performed the experiments. TNGN performed data analysis. TNGN, CLL, and JYL wrote the paper. All authors read and approved the final manuscript.

## Supplementary Figure Legends

**Supplementary Figure 1. Intron number, intron characteristics, and splicing efficiency in P. *bursaria*.**

A. The majority of *P. bursaria* genes contain multiple introns. Barplots show the distribution of the numbers of introns in *P. bursaria* genes. B. Distribution of intron lengths (in nucleotides) in *P. bursaria* (n=39,715 introns). C. *P. bursaria* introns contain conserved 5’ and 3’ splice sites. Sequence logos of the 5’ splice site and 3’ splice site were generated using ggseqlogo R. The height of each letter represents its relative frequency at that position, with letters arranged in descending order of probability. In the 5’ splice site, position 1 represents the first nucleotide of the intron. In the 3’ splice site, position-1 represents the last nucleotide of the intron. D. Most introns have a lower GC content than flanking exons. Boxplots of the GC content distribution of introns and flanking exons in *P. bursaria.* The number below each boxplot indicates the average GC content ± standard error. ***, p-value < 0.001, Wilcoxon U test.

**Supplementary Figure 2. Sequence and secondary structure of UsnRNAs in *P. bursaria*.** Conserved U1, U2, U4, U5, and U6 snRNAs in the *P. bursaria* genome. Key functional motifs are highlighted in color.

**Supplementary Figure 3. Illustration of five alternative splicing events.**

**Supplementary Figure 4. Intron retention rate (PSI) distribution and the top 30 splicing-related genes based on intron numbers with significant coefficients of differential splicing.**

A. Most introns are spliced efficiently in *P. bursaria.* Boxplots show the distribution of intron retention rate (PSI) across 16 samples collected from symbiotic and aposymbiotic cells. For each sample, the number of detected introns is indicated. B. The top 30 differentially expressed splicing-related genes that have the greatest numbers of DSIs showing significant coefficients of differential splicing.

**Supplementary Figure 5. Short intron evolution in *Tetrahymena* species.**

A. A diagram showing the phylogenetic relationships between different *Tetrahymena* species. Introns in *T. thermophila* were assigned to different age groups based on their conservation between *T. thermophila* and the other species. Age 1 group represents the youngest introns and Age 4 group represents the oldest introns. B. Young introns exhibit higher intron retention than old introns. Boxplots show the intron retention rate for each intron age group. ***, p-value < 0.001, Mann-Whitney U test. C. Young introns have a higher GC content than old introns. Boxplots show intron GC content in each intron age group. ***, p-value < 0.001, Mann-Whitney U test. D. Young introns present a wider intron length distribution than old introns. Boxplots show intron length (in basepairs) for each intron age group. ***, p-value < 0.001, Kolmogorov-Smirnov test. E. Genes containing old introns are more strongly expressed than those solely having young introns. Boxplots show expression in each gene group based on the maximum intron age in those genes.

***, p-value < 0.001; NS, p-value > 0.05, Mann-Whitney U test.

**Supplementary Tables**

**Supplementary Table 1. Intron annotation.**

**Supplementary Table 2. Splicing-related genes in different organisms.**

**Supplementary Table 3. Human splicing-related proteins specifically absent only in ciliates or yeast.**

**Supplementary Table 4. Intron retention rates of different samples.**

**Supplementary Table 5. Differentially spliced introns.**

**Supplementary Table 6. Significantly enriched GO terms in the genes containing DSIs.**

**Supplementary Table 7. Coefficients (β1) of splicing-related genes from the linear model of DSI PSI values, based on 992 DSIs and 147 differentially expressed splicing-related genes.**

