## Supplementary figures and images for "Splicing regulation and intron evolution in the short-intron ciliate model of endosymbiosis *Paramecium bursaria*"

### Supplementary Figure 1

#### Supplementary Figure 1

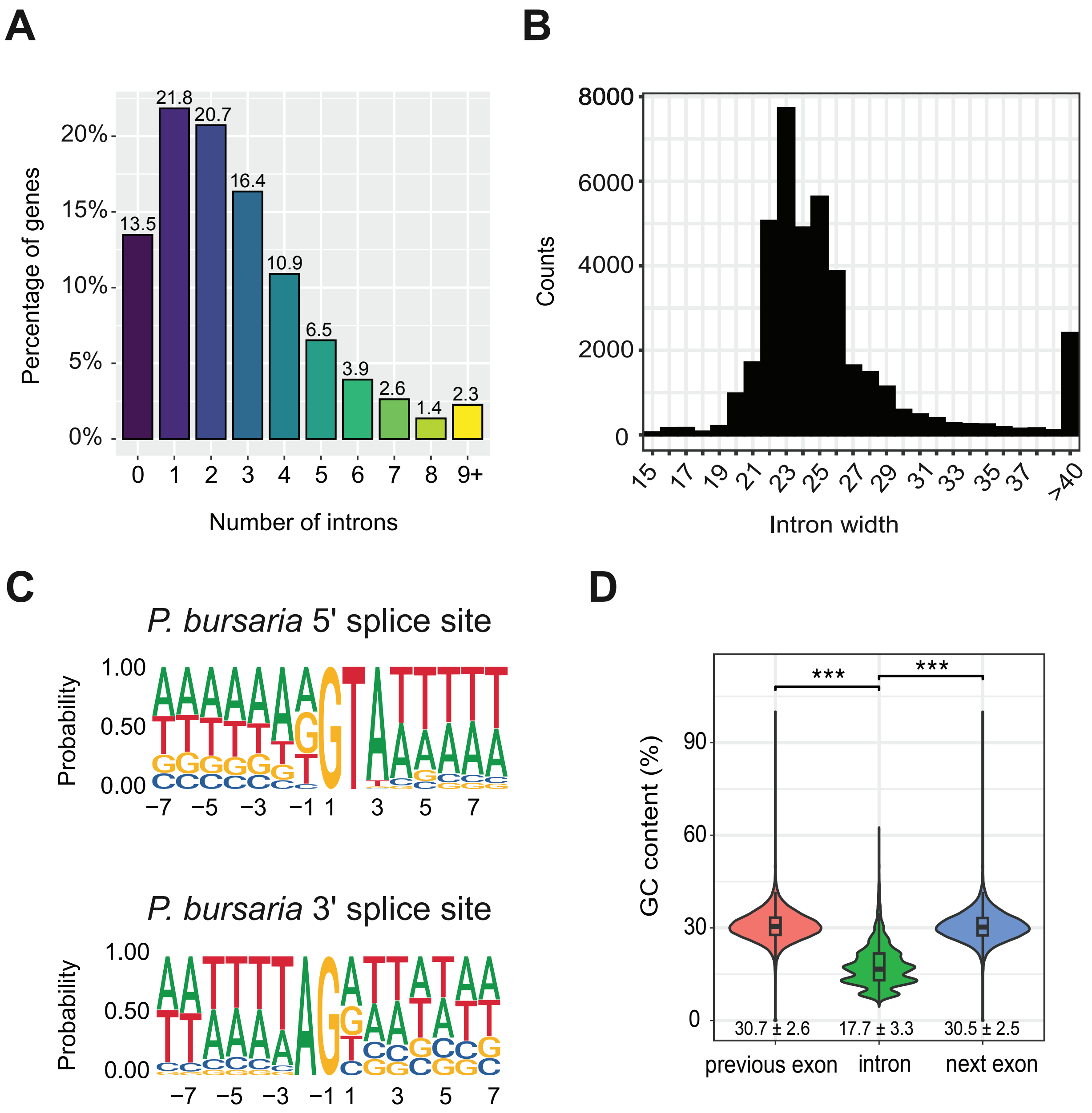

### Supplementary Figure 2

#### Supplementary Figure 2

##### U1 snRNA

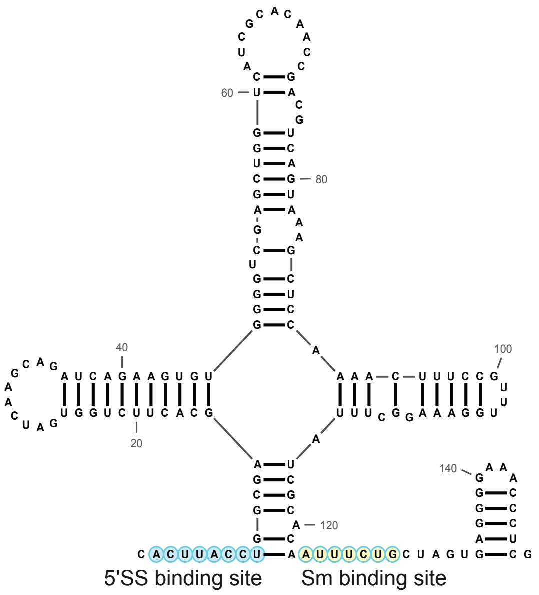


##### U5 snRNA

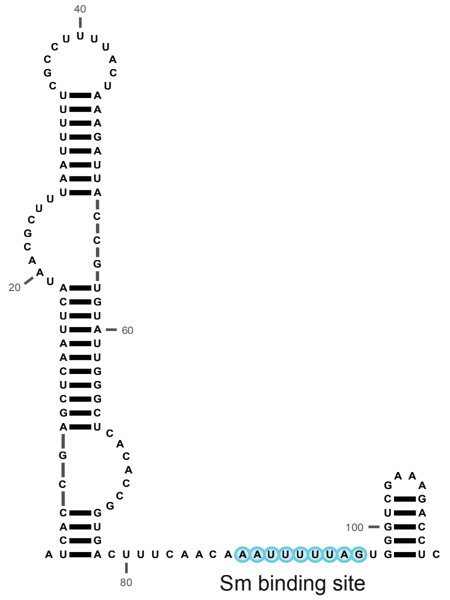


##### U2 snRNA

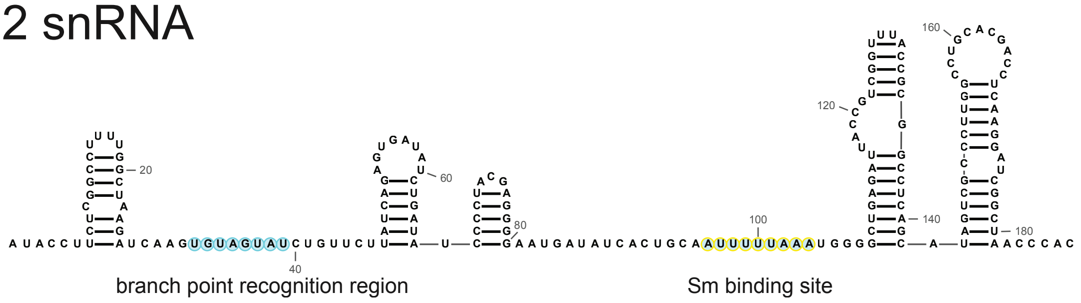


##### U4 snRNA

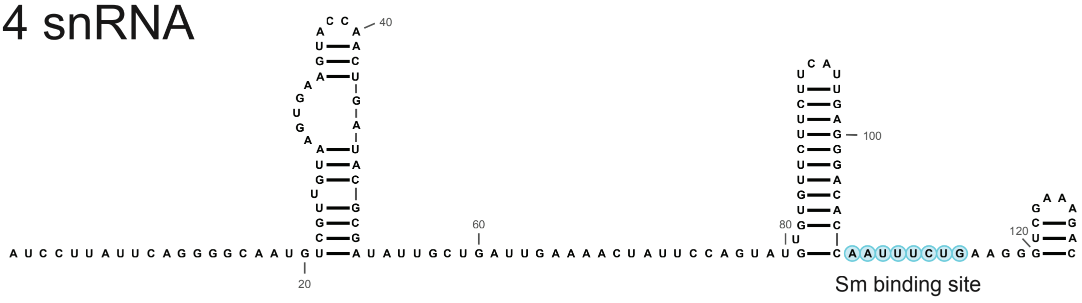


##### U6 snRNA

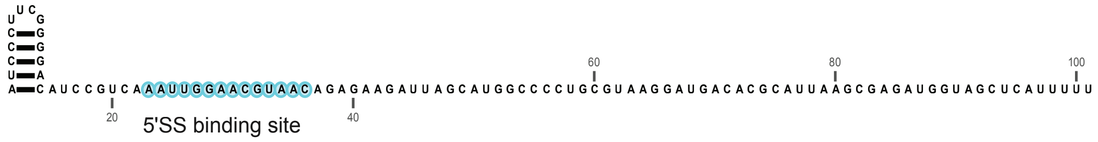

### Supplementary Figure 3

#### Supplementary Figure 3

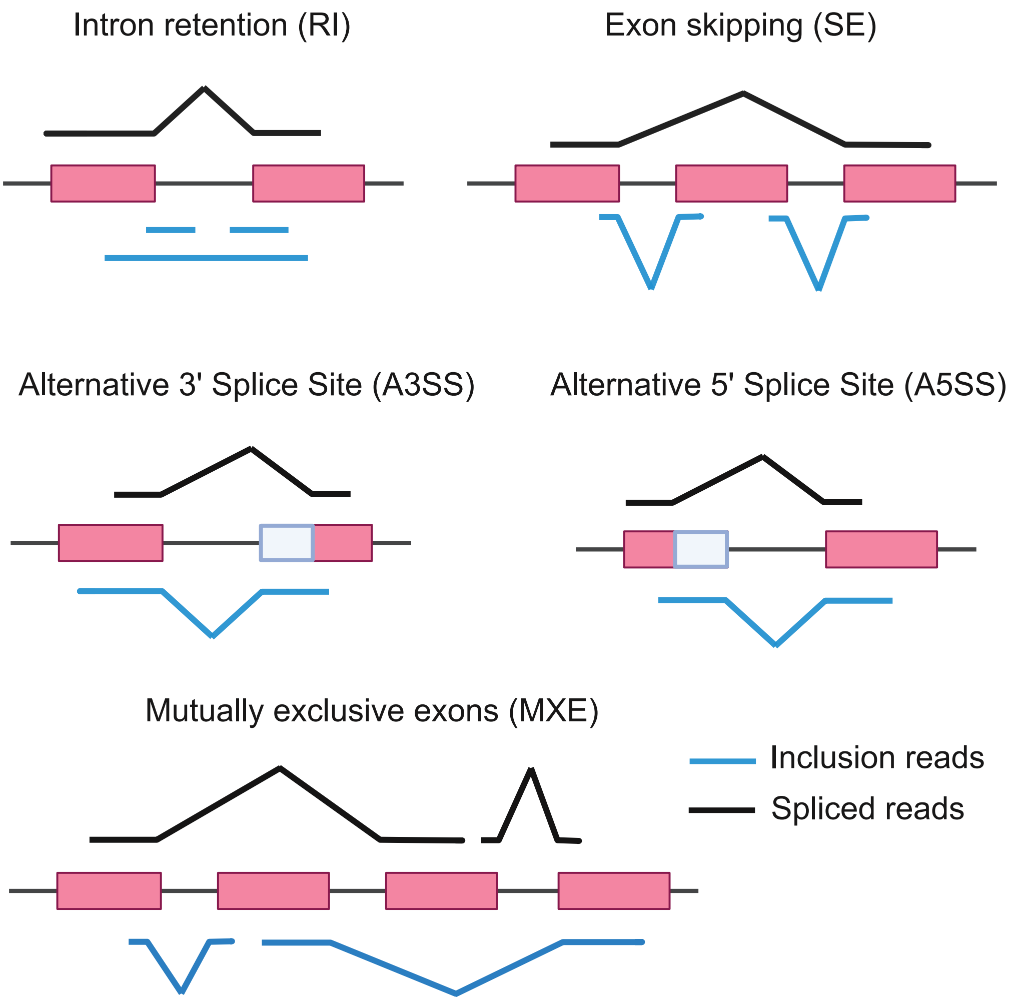

### Supplementary Figure 4

#### Supplementary Figure 4

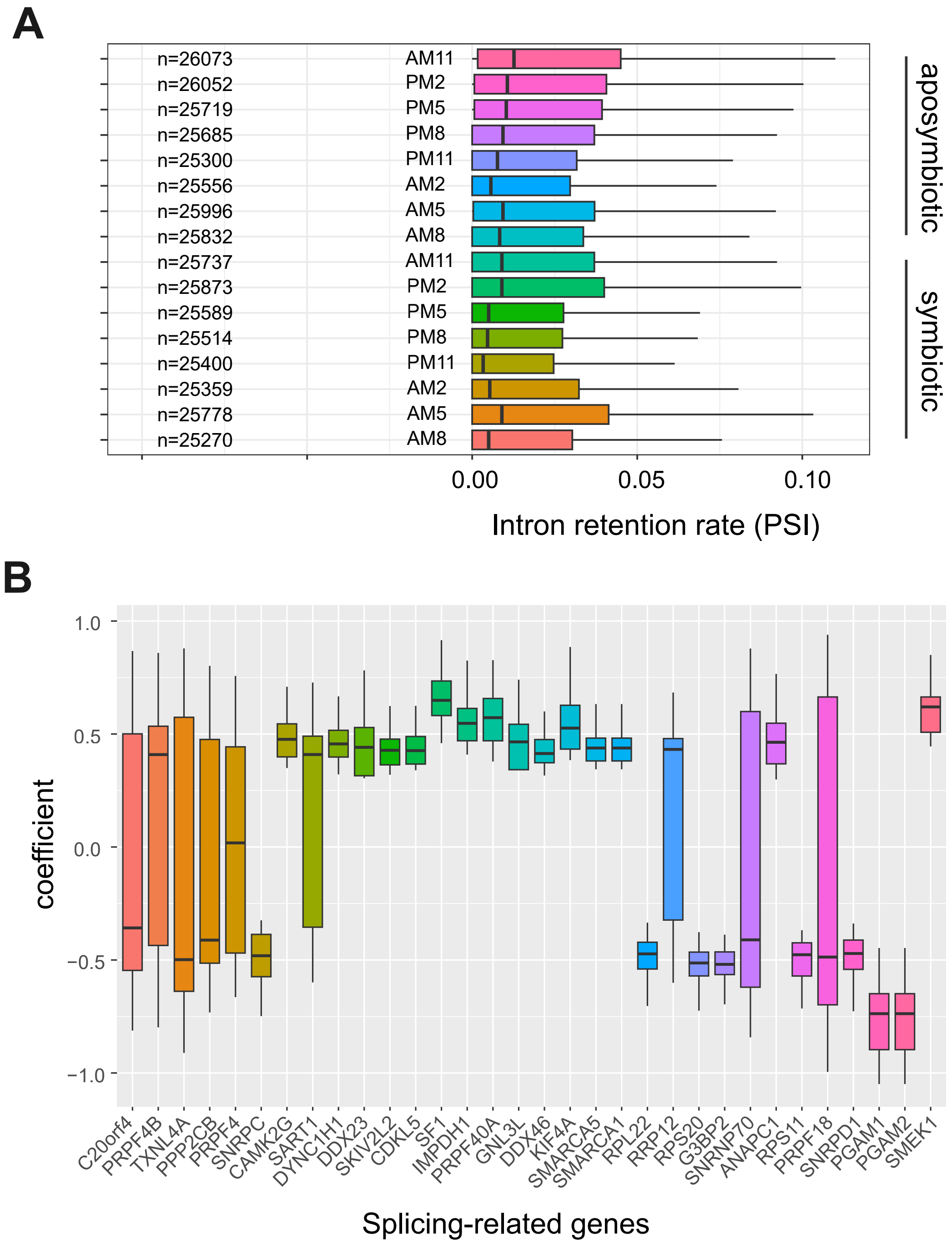

### Supplementary Figure 5

#### Supplementary Figure 5

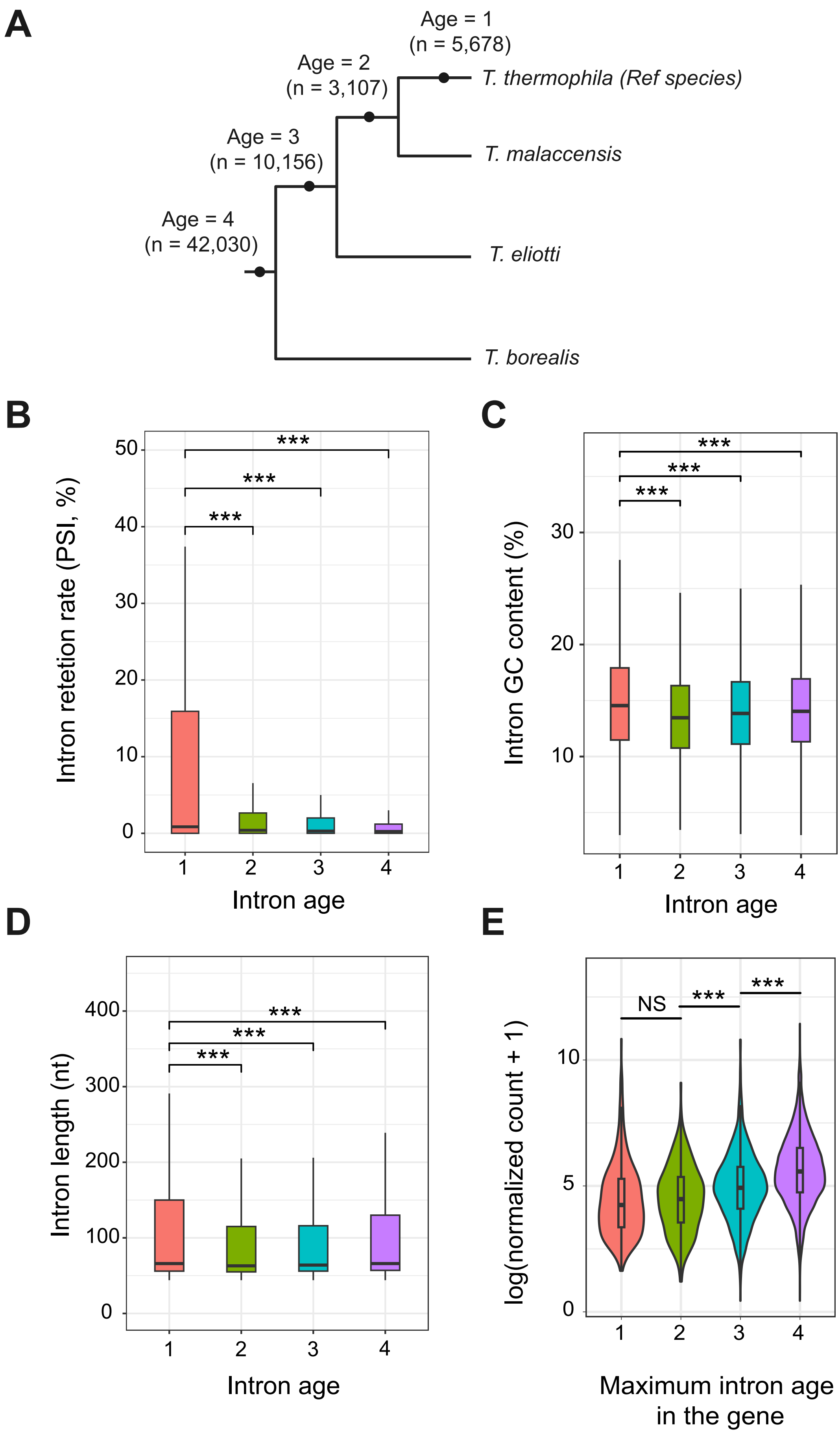
